# Molecular noise of innate immunity shapes bacteria-phage ecologies

**DOI:** 10.1101/399527

**Authors:** Jakob Ruess, Maroš Pleška, Câlin C Guet, Gašper Tkačik

## Abstract

Mathematical models have been used successfully at diverse scales of biological organization, ranging from ecology and population dynamics to stochastic reaction events occurring between individual molecules in single cells. Generally, many biological processes unfold across multiple scales, with mutations being the best studied example of how stochasticity at the molecular scale can influence outcomes at the population scale. In many other contexts, however, an analogous link between micro- and macro-scale remains elusive, primarily due to the challenges involved in setting up and analyzing multi-scale models. Here, we employ such a model to investigate how stochasticity propagates from individual biochemical reaction events in the bacterial innate immune system to the ecology of bacteria and bacterial viruses. We show analytically how the dynamics of bacterial populations are shaped by the activities of immunity-conferring enzymes in single cells and how the ecological consequences imply optimal bacterial defense strategies against viruses. Our results suggest that bacterial populations in the presence of viruses can either optimize their initial growth rate or their steady state population size, with the first strategy favoring simple and the second strategy favoring complex bacterial innate immunity.

## Introduction

One of the major challenges in biology is to understand how interactions between individual molecules shape living organisms and ultimately give rise to emergent behaviors at the level of populations or even ecosystems. At the very bottom of this hierarchy, inside single cells, interacting biomolecules such as DNA or proteins are often present in small numbers, giving rise to intrinsic stochasticity of individual reaction events [1, 2]. As a result, genetically identical organisms occupying identical environments can express different phenotypes [3, 4] and make different decisions when presented with identical environmental cues [5, 6]. This *molecular noise* is known to be the cause of biologically and medically important traits of bacteria such as persistence in response to antibiotics [7, 8] and competence during acquisition of heterologous DNA [9]. However, while its causes and consequences are relatively well-studied at the organismal level [10, 11, 12, 13], how molecular noise propagates to higher scales of biological organization to affect the ecology and evolution of organisms remains mostly unknown [4]. Recently it has been shown that ecosystems can follow surprisingly deterministic trajectories despite the prevalence of stochastic events [14, 15], yet these trajectories could themselves be strongly influenced by molecular noise. Thus, the extent to which ecological interactions are affected by molecular noise, and the extent to which these ecological consequences feed back to reshape individual traits, remain to be explored.

Perhaps the most prevalent biological systems in which molecular noise is thought to play an important role are restriction-modification (RM) systems [16]. Present in nearly all prokaryotic genomes [17], RM systems are a highly diverse class of genetic elements. They have been shown to play multiple roles in bacteria as well as archaea, including regulation of genetic flux [18] and stabilization of mobile genetic elements [19], but have most frequently been described as primitive innate immune systems due to their ability to protect their hosts from bacterial viruses [20]. When a virus (bacteriophage or phage) infects a bacterium carrying an RM system, the DNA of the phage gets cleaved with a very high probability, thus aborting the infection. With a very small probability, however, the phage can escape and become immune to restriction by that specific RM system through epigenetic modification, leading to its spread and potentially death of the whole bacterial population in absence of alternative mechanisms of phage resistance [21]. Thus, in the context of RM systems, molecular noise occurring at the level of individual bacteria can have profound ecological and evolutionary consequences. Because RM systems are ultimately based on only two very well characterized enzymatic activities (restriction and modification) [22], they represent a simple and tractable biological system in which we can investigate propagation of effects of molecular noise across different scales of biological organization.

Here, we mathematically model the action of RM systems from individual molecular events occurring inside a single cell, through individual bacteria competing in a population, to interactions between populations of bacteria and phages in a simple ecological setting, as shown in Fig 1. We demonstrate that, by imposing a tradeoff between the efficiency and cost of immunity, molecular noise in RM systems occurring at the level of individual bacteria has consequences that propagate all the way up to the ecological scale, and that the ecological consequences in turn imply the existence of optimal bacterial defense strategies against phages.

**Figure 1:**
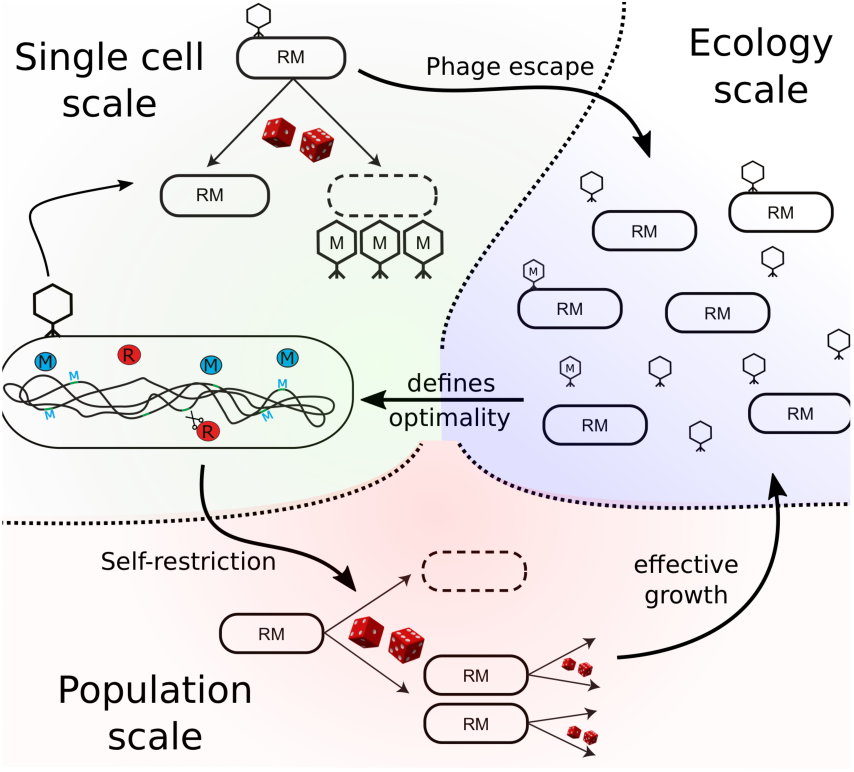
Effects of molecular noise at different scales of biological organization. In single cells (green shade), stochastic activity of restriction-modification enzymes (R - red, M - blue) may inadvertently lead to full methylation of invading phage DNA in a process known as “phage escape” from the bacterial immunity, resulting in direct consequences at the ecological scale (blue shade). Also, stochastic activity of the same enzymes may lead to accidental cutting of bacterial own DNA in a process known as “self-restriction,” resulting in lowered bacterial growth rate at the population level (red shade). Trade-offs between self-restriction and phage escape at the ecology scale in turn influence R and M enzyme activities in surviving cells, thereby optimally balancing the efficiency and cost of immunity.

## Results

### Self-restriction in single cells and in growing populations

RM systems consist of two enzymes, a restriction-endonuclease R, that recognizes and cuts specific DNA sequences (restriction sites), and a methyl-transferase M, that recognizes the same DNA sequences and ensures that only invading phage DNA can be cut by the endonuclease while the bacterial DNA remains methylated and protected. However, since chemical reactions occur stochastically, RM systems can produce errors and fully methylate invading phage DNA before it is cut and degraded (phage escape) [23]. Similarly, it is possible that newly replicated restriction sites on the bacterial DNA, which are originally unmethylated, are accidentally cleaved instead of methylated (self-restriction) [24].

Inside a single cell, the probability of such self-restriction events depends on the total activity, *r*, of all restriction endonuclease molecules R, the total activity, *m*, of all methylaze molecules M, as well as the bacterial replication rate λ, since λ determines the rate at which new unmethylated restriction sites are generated. To investigate how self-restriction depends on these parameters, we model the corresponding biochemical reactions at each individual restriction site on the bacterial DNA with the stochastic reaction network displayed in Fig 2a (see SI Appendix Section S.1). The time rs until the first self-restriction event in a given cell—i.e., until that cell’s death or substantial reduction in growth rate—can be obtained as the time when the first restriction site is cut, that is as 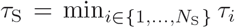 bacterial DNA with *N*_S_ restriction sites, where *τ*_*i*_, *i* = 1,…, *N*_S_ are the waiting times for cutting events at individual sites. It can be shown that all *τ*_*i*_ follow a phase-type distribution (see [25] and Fig 2b,c):

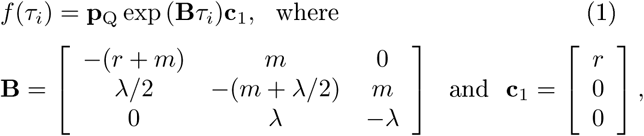

with **p**_Q_ = [*p*_0_ *p*_1_ *p*_2_] being the initial methylation configuration, i.e., the proportion of restriction sites that are unmethylated (*p*0), hemi-methylated (*p*_1_) and doubly-methylated (*p*_2_); see SI Appendix Section S.l.

**Figure 2:**
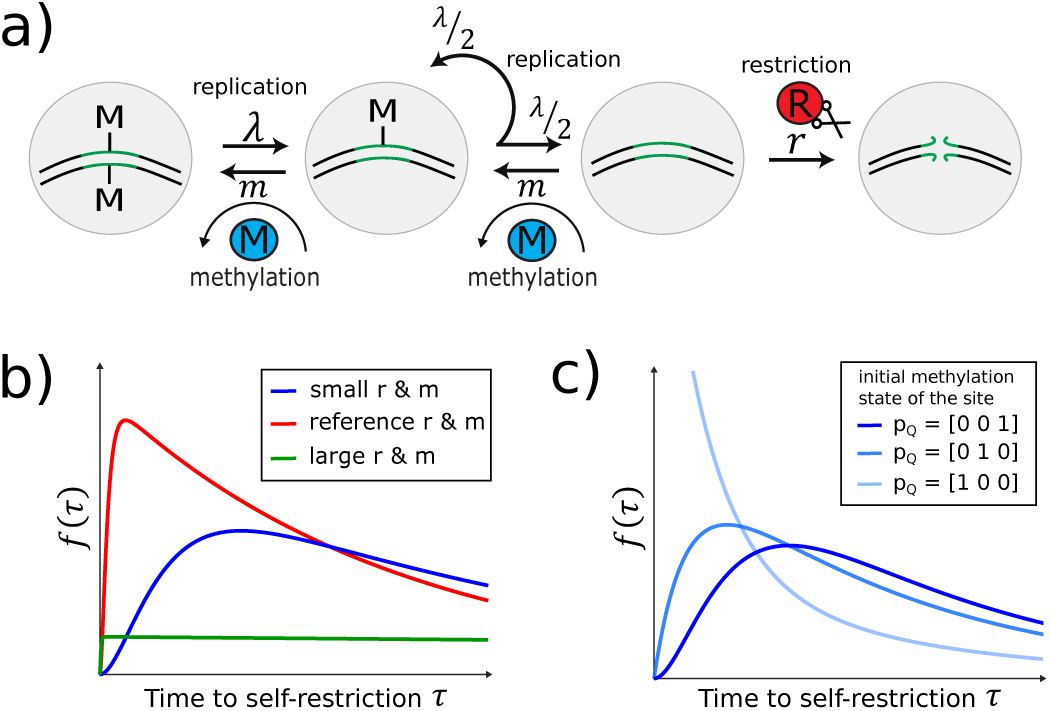
Self-restriction in single cells. **(a)** Model of the individual-site methylation dynamics. The restriction site can either be doubly-methylated (first circle), hemi-methylated (second circle), or unmethylated (third circle). Methylation events happen at a rate proportional to the activity *m* of M. During growth at rate λ, the DNA and its restriction sites are replicated, with the newly synthesized DNA having no methylation marks; for our model, growth is thus identical to demethylation reactions denoted in the figure. Restriction is assumed to be lethal (no DNA repair) and can happen when a site is unmethylated at a rate proportional to the activity *r* of R, leading to cell death (fourth circle), **(b)** Distribution of time to self restriction, *f*(τ), depends on enzyme activities, *r* and *m*, for a restriction site that is initially doubly-methylated. Increasing values of *r* lead to decreasing expected times to self-restriction for all *m* (cf. blue vs red). Given *r*, as *m* grows large, the site will almost never be unmethylated and cut, yielding a very broad distribution (green). τ = τ _ref_ = 1.7 · 10^−2^ min^−1^; reference (red) values: *m*_ref_ = 0.05 min^−1^, *r*_ref_ = 0.1 min^−1^; “Small” (“large”) *r, m* values are 2e-fold lower (e-fold higher) than reference values. **(c)** Dependence of self-restriction on the initial methylation configuration, **P**_**Q**_ (dark blue = doubly-methylated; blue = hemi-methylated; light blue = unmethylated) at *r*_ref_,*m*_ref_.

Equation [1] allows us to derive the expected time until self-restriction of a single site as

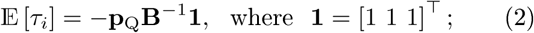

more generally, Fig 2b shows how the distribution of waiting times depends on the restriction rate ***r*** (increasing the probability of the site getting cut when it is unmethylated) and the magnitude of *m* relative to λ (which decreases the probability that the site is unmethylated in the first place).

Fig 2c shows that time to self-restriction at a single site depends essentially on an unknown quantity, the methylation configuration **p**_Q_. We will now proceed to show that when we consider an exponentially growing population of bacterial cells, the configuration **p**_Q_ can no longer be freely chosen, and has to be determined self-consistently instead. Intuitively, this is because when the bacterial population is in steady-state growth, new unmethylated sites are constantly replenished by replication, while cells with more unmethylated sites are simultaneously and preferentially being removed, as illustrated in Fig 3a and required by Eq [2]. These two forces, generation of new unmethylated sites and their preferential removal, will push any initial **p**_Q_ towards a unique steady state equilibrium.

**Figure 3:**
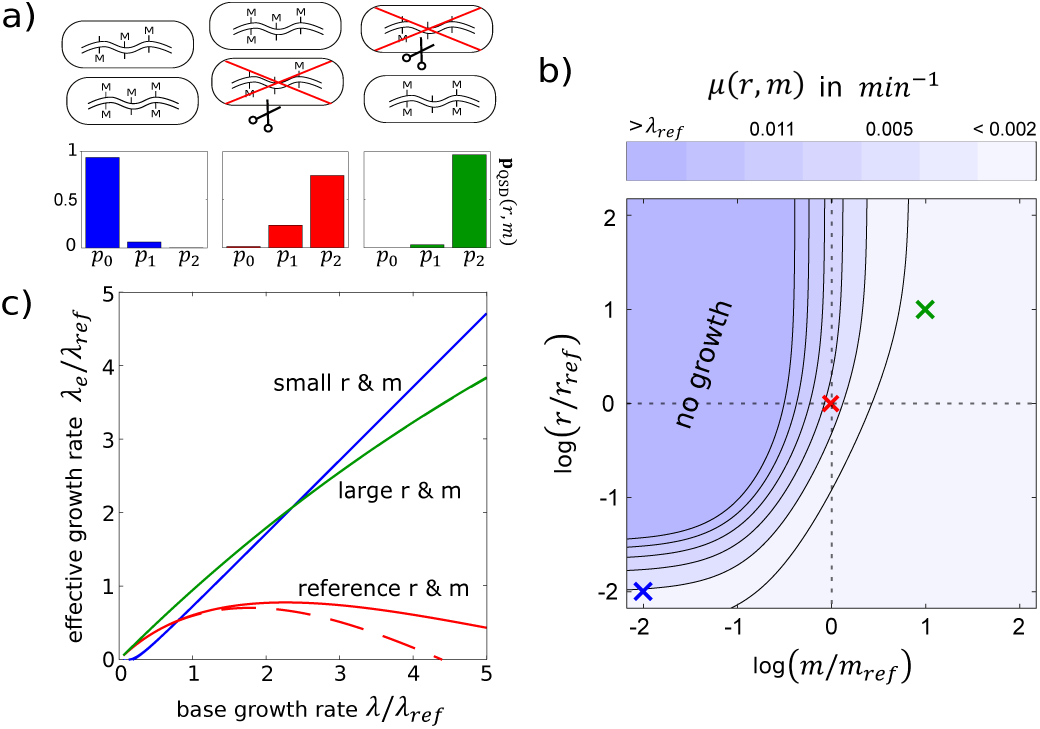
Growth of self-restricting populations, (a) Top: Selective cutting of cells (red ‘X’) with unmethylated restriction sites biases growing populations towards methylation configurations with larger numbers of methylated sites. **Bottom:** The long-term population distribution of methylation configurations, **PQSD** (*P*_*0*_ – unmethylated, *P*_1_ – hemi-methylated, *P*_2_ *-* doubly-methylated; see SI Appendix Section S.1). Increasing either *r* or *m* reduces the probability of finding unmethylated restriction sites in the population. Three example choices for *r, m* (blue, red, green) and λ _ref_ are same as in Fig 2 and are denoted with crosses in panel b and corresponding colors in panel c; *N*_*s*_ *=* 5 is chosen for illustration purposes, **(b)** Dependence of the self-restriction rate, μ (*r, m*, λ _ref_), on the enzyme activities, **(c)** Effective population growth rate λ _e_ = λ — μ (*r, m*, λ) as a function of the replication rate λ for different enzyme activities *r* and *m.*For the reference parameters (red), the self-restriction rate computed using the quasi-steady state distribution **P**_**QSD**_(solid) is significantly different when compared to an estimate based on fully equilibrated **P**_**Q**_ in which restriction does not lead to an absorbing state (dashed).

Mathematically, assuming that the methylation dynamics in all cells are equilibrated and that cells cannot be distinguished, the internal methylation configuration of any randomly chosen cell at any time during growth of the population can be derived from the quasi-stationary distribution **p**_QSD_(*r,m*) of the individual-site methylation process in Fig 2a (see SI Appendix Section S.l). **p**_QSD_(*r,m*) is the equilibrium distribution of the stochastic process conditional on it not having reached the absorbing state where the DNA is cut and the cell has died (Fig 3a); in short, methylation and growth equilibrate “*in all directions except the one leading towards self-restriction”.* Then, setting **p**_Q_ = P_QSD_(*r,m*) in Eq [1] reduces the phase-type distribution *f*(τ<u>_*<u>i</u>*_</u>) for the time τ<u>_*<u>i</u>*_</u> until self-restriction at an individual restriction site to a single exponential, implying further that the waiting time 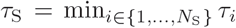 until self-restriction of any site in the cell is also exponentially distributed. Consequently, we are led to the main result of this section: growth with self-restriction can be rigorously modeled at the population level with a Markov birth-death process for which the expected population size *n*(*t*) follows a simple ordinary differential equation

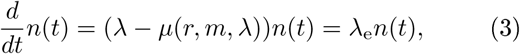

where λ_e_(*r, m*, λ) = λ — μ(*r, m*, λ) is the effective growth rate and μ(*r, m*, λ) is the rate of self-restriction, defined as the inverse of the per-cell expected waiting time until self-restriction

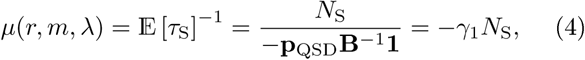

with γ_1_ being the largest eigenvalue of **B** (an explicit stochastic simulation validating this analytical result is provided in the SI Appendix Section S.2).

Equation [4] allows us to straightforwardly evaluate the reduction in the population growth rate due to random self-restriction events in single cells for any given pair of enzyme activities, ***r*** and ra. To study possible qualitative effects of self-restriction, we explore in Fig 3b a wide range of enzyme activities for a system with *N*_S_ = 5 restriction sites (chosen, for illustration purposes, significantly smaller than the typical number of sites recognized by real RM systems). We find that the main determinant of self-restriction is the activity ra of the methyl-transferase and that the effects of molecular noise can be suppressed by sufficiently increasing ra. Furthermore, so long as ra is large enough such that unmethylated restriction sites are only rarely available, μ(*r, m*, λ) lies on a large plateau of low self-restriction and changes only little with ***r*** and ra, suggesting that stochastic fluctuations in enzyme activities would only have minor consequences for the population, especially when they are positively correlated, as would be the case if R and M enzymes were expressed from the same operon (SI Appendix Section S.3).

The (*r, m*) plane in Fig 3b contains a transition region that separates the large plateau with low self-restriction from the plateau where self-restriction is severe enough to stop the population growth altogether. We have chosen our reference (red) parameter values (*r*_ref_, *m*_ref_) to lie in this transition region, and explored the regime with an e-fold higher rates (“large r & m”, indicated by green), and with 2e-fold lower rates (“small r & m”, indicated by blue) in Fig 3b, c. The comparison of these three regimes in Fig 3c is most clear when the effective growth rate is shown as a function of λ, the rate at which the cells, and thus the restriction sites, are replicated. In the “small r & m” regime, self-restriction is so infrequent that it can easily be outgrown by replication (except at very low λ). In the “large r & m” regime, *m* is sufficiently high to keep the restriction sites protected and thus self-restriction is rare, except at extremely large λ, where the green curve falls below the blue curve. In the reference regime, *r* is too large and ra not high enough to protect, so self-restriction can not be “outgrown”; effective growth thus falls significantly below λ. Our numerical analyses further show that the self-restriction rate μ(*r, m*, λ) grows faster-than-linearly with λ (SI Appendix Section S.1), causing the effective population growth to slow down and ultimately drop to zero at high enough λ.

We end this section by highlighting a non-trivial interaction between the single-cell and population-scale processes. While increasing the activity *r* of the endonuclease always decreases the effective growth rate of the population due to self-restriction, the effect can be smaller than expected from the single-cell analysis (dashed lines in Fig 3c). This is because high values of ***r*** feed back through the population scale to bias the steady-state distribution of methylation configurations away from cells with lots of unmethylated sites, as shown in Fig 3a, making self-restriction less likely. Implicit feedback effects of this type frequently give rise to complex dynamics in multi-scale models.

### Phage escape

RM systems lower the growth rate of the population due to self restriction, especially when the activity ra of the methyl-transferase is small. Upon infection by a phage, however, small values of ra are advantageous, making it less likely that the unmethylated phage DNA will get methylated and escape the immune system before it can be cut by the restriction endonuclease.

Assuming that all restriction sites are identical and independent, the probability of phage escape can be calculated [26] as

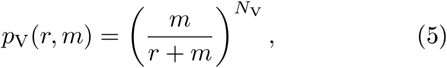

where *N*_V_ is the number of restriction sites on the phage DNA. From Eq [5] it is straightforward to see that *p*_V_ (*r, m*) is monotonically increasing in ra and decreasing in *r*. One might therefore expect that the balance between avoiding self-restriction that favors high *m*, Eq [4], and minimizing phage escape that favors low *m*, Eq [5], would impose a tradeoff and thus lead to an optimal value of ra. However, this is not the case, because the phage escape probability *p*_V_(*r, m*) and the population self-restriction rate μ(*r, m*, λ) can both approach zero so long as *r* and ra both increase to infinity but *r* does so faster. While mathematically possible, this limit is, however, not biologically relevant: large enzyme expression levels should incur a cost (metabolic or due to toxicity) for the cells [27, 28], which we sought to incorporate into our model by including a growth rate penalty proportional to the activity of restriction and methylation enzymes, i.e., λ_e_(*r, m*, λ) = μ — μ(*r, m*, λ) — *c*_*r*_*r* — *c*_*m*_*m*. Interestingly, it can be verified that our reasoning is valid only because two subsequent demethylation events need to occur to create a restriction-susceptible site on the bacterial DNA (SI Appendix Section S.1). If hemi-methylated sites could be recognized by the restriction endonuclease, or if both methyl groups could be lost in a single event, our initial expectation about the existence of the tradeoff would be correct, and a particular choice of *r* and ra values would simultaneously minimize the phage escape and self-restriction, even in the absence of the expression cost for R and M.

Our model can be generalized to multiple coexisting RM systems that recognize different restriction sites and operate in parallel, as is often observed for bacteria in the wild [17]. This provides increased protection from phages since the phage has to escape all RM systems to infect successfully. However, multiple RM systems also imply that the bacteria either have to pay higher expression and self-restriction costs or that they have to rebalance the expression levels of the enzymes such that lower self-restriction rates per RM system are obtained with the same overall enzyme activity. Allowing bacteria to have multiple RM systems, but assuming for the sake of simplicity that these systems are all equivalent in terms of enzyme activities and number of recognition sites, we obtain the phage escape probability for *k* RM systems as 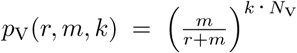, with the corresponding growth rate being

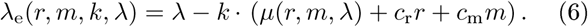

### Population dynamics in the presence of phages

What is the combined effect of phage escape and self-restriction in simple bacteria-phage ecologies? To investigate this question, we studied several different ecological contexts analytically as well as numerically (see SI Appendix Section S.4) and performed an in-depth exploration of a scenario where bacteria are trying to colonize environments in which lytic phages are already present in large numbers.

### Bacterial population dynamics with constant phage load

To set the stage, we first considered a minimal, mathematically simple but biologically unrealistic scenario where phage escape is modeled deterministically and the number of phages in the environment is constant in time. Typically, this will not be the case, as phages will not only kill bacteria upon successful infection and phage escape, but will also increase in their number. Nevertheless, we start with this setup that can be mathematically understood in its entirety, and relax its crucial assumptions later.

The dynamics of the bacterial population in the presence of obligatorily lytic phage, *n*(*t*), can be written as follows:

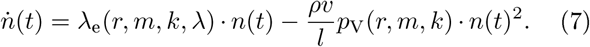

The first term on the right-hand side describes the growth of the population at an effective rate of μ _e_ (*r, m, k*, λ) that already accounts for self-restriction. The second term models phage escape as a deterministic decrease in bacterial population size. Here, *v* is the phage population size which we have taken to be a constant parameter of the environment; phages enter bacterial cells at a rate proportional to *vn*(*t*), as prescribed by mass-action kinetics, with a proportionality constant given by the phage adsorption rate, *ρ.* The rate of successful infections is then given by *pvp*_V_(*r,m,k*)*n*(*t*). We assume that each successful infection event wipes out a fraction, 0 < *1/l* < 1, of the total population (i.e., on average, *n(t)/l* bacteria die following phage escape), yielding the full Eq [7].

The growth dynamics has one biologically-relevant fixed point that represents the steady-state bacterial population size:

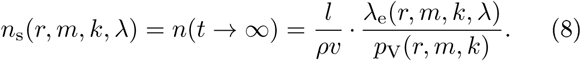

In this minimal-model scenario, which is denoted as (o) in Table 1, ***n*_*s*_** quantifies the bacterial performance in terms of long-term population size (see SI Appendix Figure S.6). The unknown parameters ρ, ***v***, and *l*, enter Eq [8] as multiplicative constants that are independent of single cell parameters. Optimal performance, which we will explore after we introduce alternative ecological scenarios below, would therefore imply finding singlecell parameters *r, m*, and ***k***, that maximize the ratio λ _e_(*r, m, k*, λ)/*p*_v_(*r,m,k*).

**Table 1:** Efficiency criteria for bacterial populations in different ecological scenarios. Minimal, constant phage load scenario, (o), and bacteria colonizing new phage-dominated environments, (i), (ii), or (iii). Optimality criterion in the first column is captured by the corresponding mathematical expression for the bacteria-phage ecology in the second column, which implies the maximization of the “bacterial performance” quantity in the third column. *c*_mut_ is the mutation rate per unit time per bacterial cell to gain resistance against phage (and 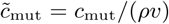) is the normalized mutation rate); τ_mut_ denotes the (random) time at which the first mutation happens.

| efficiency criterion | mathematical expression | bacterial performance | summary figure |
| --- | --- | --- | --- |
| (o) long-term population size in constant phage load scenario | $n(t \rightarrow \infty)$ | $\lambda_e/p_V$ | Fig 4a |
| (i) expected growth until phage escape | $\mathbb{E}[n(\tau_p)] - n_0$ | $\lambda_e/p_V$ | Fig 4a |
| (ii) prob. that mutation happens before phage escape | $\mathbb{P}(\tau_{\text{mut}} < \tau_p)$ | $\tilde{c}_{\text{mut}}/(\tilde{c}_{\text{mut}} + p_V)$ | Fig S9a |
| (iii) expected integrated population mutation rate | $\mathbb{E}[\int_0^{\tau_p} c_{\text{mut}} n(t) dt]$ | $\tilde{c}_{\text{mut}}/p_V$ | Fig S9b |

### Bacterial population dynamics in ecologically more realistic scenarios

While the minimal model is mathematically convenient, the fact that it doesn’t include phage population size as a dynamical variable and that it abstracts the appearance of methylated phage with a single deterministic term in Eq [7] is clearly an oversimplified representation of phage ecology. To obtain biologically more meaningful insights, we therefore have two options: we can either consider a more realistic model that explicitly tracks phage dynamics or we can represent phage escape events stochastically and avoid having to explicitly model methylated phage by studying bacterial growth only until the (random) time at which phage escape happens for the first time.

To explore the the first option, we extended an established deterministic model of bacteria-phage ecology [29] to track the population dynamics of bacteria with and without RM systems and both susceptible and methylated phages (see SI Appendix Section S.4.2). By numerically integrating this population model for more than a million parameter combinations for the activity of restriction (*r*) and methylation (*m)* enzymes, we find that whether or not phages will ultimately take over the population depends on the ecological parameters (e.g. phage adsorption rate, rate of spontaneous phage inactivation, etc.) but is completely independent of RM system efficiency. While this result clearly depends on the fact that phages cannot go extinct in ecology models based on ordinary differential equations, it nevertheless suggests that long term bacterial population sizes, such as the steady state, *n*_s_, of the minimal model, (o), cannot be a biologically realistic measure for the efficiency of RM systems and that the task of RM systems cannot be to prevent phage escape but only to delay it as much as possible to give bacteria enough time to develop alternative mechanisms of phage resistance through genetic mutations [21]. We conclude that the relevant option to study is therefore the second one: how RM systems impact bacterial growth until the first phage escape event.

To explore the second option, we formulated several efficiency measures that quantify how RM systems can help bacterial populations before the first phage escape event:

i. How much can a bacterial population grow before the first phage escape event happens?
ii. What is the probability that an immunity conferring mutation happens in any bacterium before the first phage escapes?
iii. What is the total number of mutations accumulated in the population when methylated phages start to spread?

Here we will show that questions (i)-(iii) can be answered rigorously if we assume that the size of the phage population remains approximately constant until the first phage escape event. An example of an ecological scenario where this assumption is realistic is that of bacteria colonizing a phage-dominated environment in which the number of phages is much larger than the number of bacteria such that the reduction in the phage population size due to unsuccessful infections is negligible. More generally, any ecological scenario in which the phage population size is for some reason in equilibrium at least until the first phage escapes on a bacterium carrying a RM system, will fulfill this assumption. Surprisingly, the results that we will find will connect back to the simple model summarized by Eq [7].

Mathematically, we consider a bacterial population of initial size *n*_0_ trying to colonize an environment containing a phage population of size ***v.*** As we have shown before, the bacterial population will initially grow exponentially at rate λ_e_ until the time τ_*p*_ at which the first phage escape event occurs. Interpreting these events as random, the crucial unknown is therefore τ_*p*_, the random time to first phage escape, characterized by its probability distribution, *f* (τ_*p*_), which we find to be given by (see SI Appendix Section S.4.3):

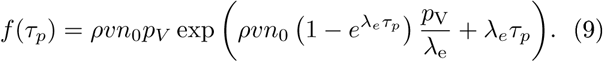

The waiting time distribution until first phage escape, *f* (τ_*p*_), allows us to analytically answer questions (i)-(iii), as summarized in Table 1 (see SI Appendix Section S.4.3). Each efficiency criterion corresponds to a maximization of the corresponding “bacterial performance” metric. By examining these metrics, we make three important observations:

First, expressions for bacterial performance in Table 1 are functions of λ_e_ and *p*_*V*_, which depend solely on the restriction rate *r*, the methylation rate *m*, and the number of concurrently active RM systems, ***k.*** This means that optimal bacterial strategies at the ecological level can be found mathematically—and possibly tuned evolutionarily—by adjusting the three parameters, *r, m*, and *k*, defined at the single-cell level.

Second, the performance of the bacterial population is independent of the initial population size, ***n*_0_**. As a consequence, there exists a unique best defense strategy against phages that is constant in time: if phage escape has not happened until a certain time during which the bacterial population has grown to a new size, the same defense strategy continues to be optimal with the initial size taken to be the new size, with no need to re-balance the activity levels ***r*** and *m* of the RM enzymes, or the number of RM systems, ***k.*** For cases (ii) and (iii), we further observe that the results are independent of the effective growth rate λ_e_. Faster growth leads to quicker increases in the probability that immunity conferring mutations happen but this is exactly compensated by the increase in probability of a phage escape event.

Third, while maximizing the growth of the bacterial population until the first phage escape event, Table 1 (i), is biologically very different from maximizing the steady-state population size under constant viral load, Table 1(o), both questions lead to mathematically equivalent problems of maximizing the ratio λ_e_/*p*_v_ with respect to single cell parameters *r, m*, and ***k.*** In the following, we focus an in-depth study of the consequences and implications of case (i). Questions (ii) and (iii) are treated in the SI Appendix (Section S.4.4).

### Tradeoffs and optimality in bacterial immunity

Can bacteria tune the single-cell parameters over evolutionary timescales in order to minimize the cost of RM systems, i.e., maximize the growth rate λ_e_, while also maximizing their efficiency, i.e., maximize λ _e_/.*P*_*v*_? Equation [6] and Table 1 assert that cost and efficiency are necessarily in a tradeoff and cannot be optimized simultaneously. This tradeoff is the first key result of the section. With no single optimum possible, we look instead for Pareto-optimal parameter combinations, (*r, m,k*), i.e., solutions for which λ_e_ cannot be further increased without reducing λ**_*e*_/*p*_*v*_** and vice versa [30, 31]. Different Pareto-optimal solutions trace out a “front” in the plot of λ_e_ vs λ_*e*_*/p*_*v*_ in Fig 4a that jointly maximizes growth rates and population sizes to the extent possible. Points in the interior of the front are sub-optimal and could be improved by adjusting parameter values, while points beyond the front are inaccessible to any bacterial population. Which Pareto-optimal solution ultimately emerges as an evolutionary stable strategy depends on the actual bacterial and phage species considered as well as their biological context. Rather than focusing on specific examples, we next establish several general results of our analysis, contrasting in particular “fast growth” bacterial strategies that maximize λ_e_ with “large size” strategies that maximize λ_e_/*p*_*v*_.

**Figure 4:**
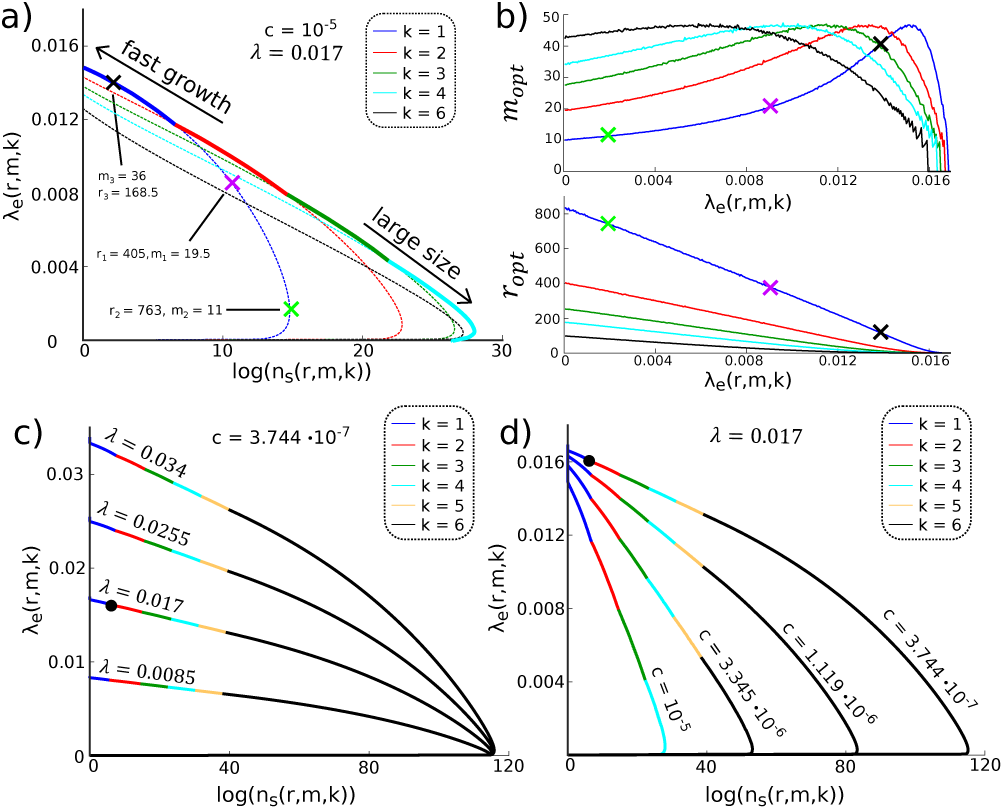
Optimal tradeoffs in the presence of phages. For all panels, we set *N*_*s*_ = 599, *N*_*v*_ = 5, and λ = 0.017 min^−1^ in line with the strain used in [24] carrying the EcoRI system, and without loss of generality we fix *l/ρ v =* lmin such that *n*_*s*_ and *λ*_*e*_*/P*_*v*_ are the same up to units, **(a)** Pareto front (thick line) for expected growth until phage escape, n_s_(*r,m,k*), and effective growth rate, λ _e_(*r,m,k*), traced out by varying *m, r*, and the number of RM systems, *k* (different colors; dotted lines show individual Pareto fronts at fixed *k*)*’, c* = *c*_*r*_ = *c*_*m*_ = 10^−5^. Examples with specific parameter values are marked on *k* = 1 front with crosses, **(b)** Optimal enzyme activities, *r*_opt_ and *m*_opt_ (in min^−1^), on Pareto fronts for different *k* (color as in a), as a function of effective growth rate, λ _e_. **(c)** Pareto fronts for different replication rates, λ, and cost *c* = *c*_*r*_ = *c*_*m*_ ≈ 3.7 · 10^−7^ chosen to make *E. coli* in [24] lie on the front (black dot, also in panel d); only *k* ≤ 6 are considered. (d) Pareto fronts for different cost *c = c*_r_ = *c*_m_ at fixed λ = 0.017 min^−1^.

We start by examining in Fig 4b the optimal enzyme activities, *m*_*opt*_ and r_op*t*_, along the Pareto fronts. For the “large size” regime at low λ_e_, the bacterial population primarily needs to defend against phage escape, favoring low *m* and high *r*, even at the cost of self-restriction. As we move towards the “fast growth” regime, ***r*** can drop to decrease the cost, but *m* must increase to protect against self-restriction, until maximal *m*_op*t*_ is reached. For even higher λ_e_, it is optimal to “shut down” the RM systems altogether to save on the cost, by tuning ***r*** and *m* simultaneously to zero. Numerical analysis (SI Appendix S.4.5) reveals that along the Pareto front of Fig 4a, the *total* cost of running the RM systems varies in precise inverse linear relationship with λ_e_. Pareto-optimal solutions are further characterized by the fact that the reduction in growth rate, λ — λ_e_, is split equally between the cost of running RM systems, *c*(***r*** + *m*), and self-restriction. If this were not the case and the cost were larger (or smaller) than self-restriction cost to growth, cells could always down- (or up-)regulate the RM system activity to trade cost for self-restriction and obtain an overall smaller total growth reduction. This universal equality of cost of running RM systems and self-restriction at optimality is the second key result of the section.

A detailed examination of the Pareto front in Fig 4a reveals a striking shift in the structure of optimal solutions as we move from “fast growth” to “large size” regime. In situations where fast growth is favored, we observe that a single RM system (*k* = 1) is optimal. In contrast, large steady-state bacterial population sizes favor *k*_opt_ > 1 RM systems, with the optimal number, *k*_opt_, set by the costs, *c*_m_ and *c*_r_, of operating the RM systems. These results are quantitatively robust to changes in replication rate, λ, as shown in Fig 4c, where Pareto fronts for different λ are nearly rescaled versions of each other. These results are also qualitatively robust to changes in the cost *c* = *c*_r_ = *c*_m_ so long as the cost is nonzero, as shown in Fig 4d.

Establishing that “fast growth” regime favors simple innate immunity with a single RM system while “large size” regime favors complex innate immunity with multiple RM systems is the third key result of this section. This result can be understood intuitively by considering under what conditions, if any, multiple RM systems could be optimal at “fast growth”. If costs for R and M enzymes are vanishingly small, a single RM system can provide arbitrarily good protection, as we showed previously. If the costs are not vanishingly small, multiple RM systems must be more costly than a single system at comparable phage escape and self-restriction rates: to keep self-restriction constant with *k* RM systems, not only does the cell require *k* times more M molecules than at *k* = 1, but their individual activities need to be higher as well, leading to a higher cost for M and thus a lower effective growth rate; thus, *k* > 1 cannot be optimal for “fast growth” and that can only be tolerated in the “large size” regime where protection from phages is more important than fast growth.

Lastly, we sought to put our results into perspective by relating them to a typical *E. coli* strain. Recent measurements [24] quantified the self-restriction rate in a bacterial population with the EcoRI system replicating at λ = 0.017 min^−1^ to be around μ ≈ 10^−3^ min^−1^. The cost of RM systems was not detectable in WT strain but could be detected in strains overexpressing M enzymes. Treating the cost ***c*** as unknown and assuming that *E. coli* is Pareto-optimal in light of criterion (i) in Table 1 (black dots in Fig 4c,d), would lead us to predict the following parameter values for the RM systems: cost *c* ≈ 3.7 · 10^−7^, enzyme activities *r* ≈ 1.2 · 10^3^ min^−1^, *m* ≈ 1.5 · 10^2^ min^−1^, with the optimal number of RM systems being at the boundary between *k* = 1 and *k* = 2. Clearly, this prediction depends on the chosen measure of the efficiency of RM systems, which is determined by the considered ecological scenario and the particular objective that bacteria have in this scenario. Consequently, the concrete numbers presented here should not be understood as general results, but rather as a demonstration of how our framework can be used to calculate optimal bacterial strategies given different modeling assumptions about the phage-bacteria ecology.

## Discussion

Despite the ubiquity of RM systems in prokaryotic genomes [17], basic ecological and evolutionary aspects of these otherwise simple genetic elements are poorly understood [20]. Although RM systems have been discovered more than six decades ago due to their ability to protect bacteria from phage [32] and this is often assumed to be their main function [33], only a few experimental studies focused on the ecological and evolutionary dynamics of interactions between RM systems and phage [34, 35]. Similarly, effects of RM systems on their host bacteria, such as their cost in individual bacteria due to self-restriction, began to be addressed quantitatively only recently [24, 36]. In this work, we bridged these two scales using mathematical modeling. Our model captures the stochastic nature of RM systems originating at the level of interacting molecules in individual bacteria and extends it all the way to the dynamics of interactions between bacterial and phage populations.

Using this approach, we analytically described tradeoffs between the cost and the efficiency in different ecological contexts of immunity conferred by RM systems. The existence of such tradeoffs was previously indicated by quantitative single-cell experiments with two RM systems isolated from Escherichia coli [24]. We used our mathematical framework to quantify these tradeoffs and to study their ecological consequences, as well as the implications that these consequences have for optimally tuning the R and M enzymatic activities at the level of single cells. Our results for different ecological scenarios suggest that we should expect observed expression levels and enzymatic activities of naturally occurring RM systems to represent adaptations to specific environmental pressures. Such “tuning” of expression levels towards optimality has previously been directly experimentally shown in different molecular systems [27]. The expression levels of both R and M should be readily tunable by mutations in the often complex gene-regulatory regions [37].

With optimal bacterial defense strategies depending on the ecological scenario and the particular objective of the bacteria (see SI Appendix Section S.4 and Table 1), making general predictions on R and M expression levels or numbers of concurrently active RM systems that we should expect to find in bacteria in the wild is difficult. However, we want to highlight that, in a given context, assuming optimality of the bacterial defense strategy allows one to make clear and quantitative predictions about the reaction rates and the number of RM systems, and im-proving these predictions to take into account more relevant biological detail (if needed and known) remains only a technical, rather than conceptual challenge. Second, for the scenario of bacteria colonizing phage dominated environments that we investigated in this paper, parameter values measured for an *E. coli* RM system put optimal solutions into a regime that permits a large variation in the optimal number of RM systems, between one to six, with relatively small changes in the effective growth rate. This observation allows us to advance the following hypothesis: the number of RMsystems in different bacterial strains and species is not a historical contingency, but an evolutionary adaptation to different ecological niches. In other words, the tradeoff between the cost and the efficiency of immunity can be partially alleviated in bacteria employing multiple RM systems. It is therefore interesting to note that many bacterial species carry multiple RM systems and the number of RM systems varies significantly among bacteria with different genome sizes and lifestyles [17, 16]. Our results indicate that different numbers of RM systems would be optimal in populations under different selection pressures (phage predation /resource limitation).

The analytical model presented here makes several simplifying assumptions. First, we consider only interactions between a single species of bacteria and a single species of phage. In natural environments, many bacterial and phage species interact and this diversity will certainly impact the resulting ecological end evolutionary dynamics [38, 39, 34, 40]. Second, we assumed the key parameters such as the numbers of restriction sites in bacterial and phage genomes to be constant in time and thus disregarded the long-term evolutionary dynamics. Bioinformatic studies have shown that many bacteria and phage avoid using restriction sites in their genomes [41, 42, 43]. Restriction site avoidance can represent an adaptive mechanism for increasing the likelihood of escape in phages [42, 44] and decreasing the likelihood of self-restriction in bacteria [45, 24]. The stochastic nature of RM systems observed at the level of individual cells is thus likely to critically shape the ecological and evolutionary dynamics of interactions between bacteria, RM systems and phage.

## Acknowledgements

The authors would like to thank Moritz Lang, Gregory Batt, Eugenio Cinquemani and Christoph Zechner for helpful discussions. JR acknowledges support from the Agence Nationale de la Recherche (ANR) under Grants No. ANR-16-CE33-0018 (MEMIP), ANR-16-CE12-444 0025 (COGEX), and ANR-18-CE91-0002-01 (CyberCircuits). CCG was supported by the HFSP Young Investigators’ grant. MP was a recipient of a DOC Fellowship of the Austrian Academy of Science at the Institute of Science and Technology Austria. GT was supported in part by the Austrian Science Fund grant FWF P28844.

